# Somatic DNA methylation heterogeneity predicts extreme transgenerational epimutation hotspots in Arabidopsis

**DOI:** 10.64898/2026.04.04.716457

**Authors:** B Thanh Vo, P Wolf, J Kim, Z Zhang, V Ramirez, B Poppenberger, K Schneitz, C Becker, M List, F Johannes

## Abstract

Spontaneous epimutations are stochastic and heritable changes in cytosine methylation that arise independently of DNA sequence alterations. In plants, they occur predominantly at CG sites, accumulate across generations at high and clock-like rates, and contribute substantially to constitutive epigenetic diversity. Although these features are well established, the developmental origin of spontaneous epimutations remains poorly understood. One model proposes that they arise during somatic growth in different cell layers of the shoot apical meristem, where lineage bottlenecks enable nascent epimutations to clonally propagate into developing organs. Such layer-specific dynamics are expected to generate spatial mosaics of methylation states within tissues that manifest as heterogeneity in bulk bisulfite sequencing data. Here, we quantified genome-wide CG methylation heterogeneity in single Arabidopsis leaves using read-level methylation discordance metrics adopted from cancer epigenomics. We identified thousands of highly heterogeneous loci and show that they overlap extreme transgenerational epimutation hotspots. Analysis of multiple leaves from the same individuals further revealed that methylation divergence at these loci scales with developmental distance and recapitulates the branching architecture of the shoot. Together, these results support a meristematic origin of spontaneous epimutations and highlight a shared susceptibility to methylation-maintenance errors in both developmental and transgenerational contexts.

---

Spontaneous epimutations are stochastic and heritable changes in cytosine DNA methylation states that arise in the absence of underlying DNA sequence alterations (1). These events occur predominantly at CG dinucleotides and are widely attributed to imperfect maintenance of methylation by DNA methyltransferases during cell divisions (1,2). Importantly, in plants, these variants accumulate across generations in a manner analogous to de novo nucleotide substitutions. Multi-generational experiments show that spontaneous epimutations occur at rates several orders of magnitude higher than DNA sequence mutations per unit time (3), accumulate in a largely neutral and clock-like fashion (4), and yield robust phylogenetic signal over short evolutionary timescales where genetic divergence is limited (5,6). The accumulation of epimutations is not uniformly distributed across the genome, but exhibits strong compartment-specific biases (3,7), including markedly elevated transition frequencies within specific classes of genes (8). In *A. thaliana*, these genomic biases correlate with CG methylation diversity patterns among natural accessions, suggesting that epimutational processes are a major source of constitutive CG methylation variation in wild populations (3,7,9,10).

Despite these insights, the developmental origin of spontaneous epimutations within individual plants remains poorly understood. One hypothesis places their formation during gametogenesis or early embryogenesis, when methylation patterns are highly dynamic (11,12), undergo genome-wide reinforcement (13), and may be vulnerable to off-target enzymatic activity (1). An alternative hypothesis proposes that spontaneous epimutations arise gradually during somatic growth (4,14), particularly in stem cells located in the central zone of the shoot apical meristem (SAM). These cells comprise a small, slow-dividing stem-cell pool that is maintained over time (15). Divisions leave one daughter in the stem-cell niche, while the other is displaced toward the SAM periphery, where it is incorporated into differentiating tissues (15,16). Epimutations arising in these stem cells can thus accumulate through self-renewal and be propagated into the lineages that form new tissues during growth. A direct test of this SAM-origin hypothesis requires time-resolved, single-cell methylome profiling of individual meristems combined with cell lineage tracing (17), which remains technically challenging (18). Nonetheless, indirect evidence from long-lived perennials strongly supports this model, because epimutation burden increases with organismal age (14), is higher in contexts of rapid growth and greater somatic cell production (19), and tracks branching topology within individuals (20).

Because all above-ground plant organs arise from a small number of precursors derived from stem cell lineages (15), newly formed epimutations can rapidly fix within specific meristematic layers (L1, L2, and L3) through drift (4,17). This layer-specific fixation is expected to produce persistent spatial mosaics of DNA methylation within organs (e.g., within fruits or leaves) and across the shoot system (**Fig. 1A**). Comparable patterns have recently been reported for somatic DNA sequence mutations in apricot and potato, where mutation profiles differed substantially between L1, L2, and L3-derived tissue layers (21,22). In bulk bisulfite sequencing of intact organs (e.g. whole leaves, or fruits) these mosaics are expected to manifest as localized DNA methylation heterogeneity at affected loci (23), because bulk methylation estimates reflect the mixture of read-level methylation states contributed by different SAM-derived tissue layers (**Fig. 1A**). Analytical approaches from cancer epigenomics have quantified this cellular heterogeneity by measuring read-level methylation discordance among neighboring CG sites on homologous DNA molecules using short-range epihaplotype information (24–29) (**Fig. 1B**). These metrics provide a tractable proxy for somatic epigenetic diversity without explicit cell-type resolution and have been used successfully to infer subclonal structure in bulk tumor samples (24).

**Figure 1:**
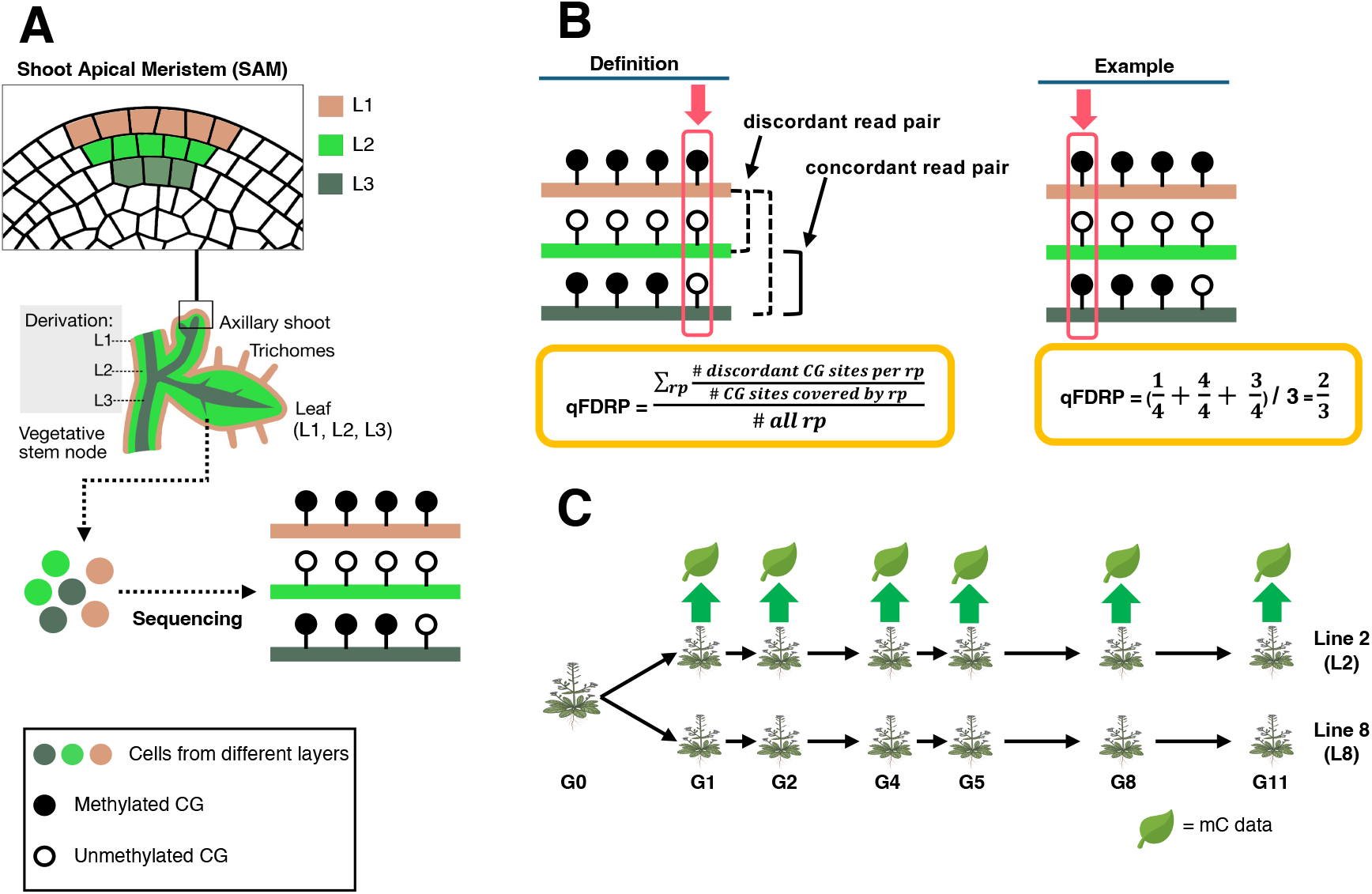
Conceptual framework and experimental design for quantifying somatic DNA methylation heterogeneity using qFDRP. A. Somatic epimutations arising in different layers of the SAM stem cells generate methylation mosaics in plant organs. B. Quantification of methylation heterogeneity using the qFDRP score. The example illustrates how discordant and concordant reads contribute to the qFDRP calculation. C. MA lines are derived from a single inbred founder plant at generation G0 and propagated for many generations (11 generations and two lines, L1 and L2, are shown here). Leaf samples from different generations (G0 – G11) were sequenced to generate WGBS data.

Here, we applied this analytical framework to map genome-wide CG methylation heterogeneity in *A. thaliana* leaves and identify loci that are particularly susceptible to somatic epimutations. Our analysis revealed that regions with the highest somatic heterogeneity within leaves overlap with extreme transgenerational epimutation hotspots, revealing a shared vulnerability to methylation maintenance errors in both developmental and transgenerational contexts. Moreover, we found that CG methylation divergence among leaves from the same plant reflected their spatial and temporal positions along the shoot axis, producing patterns that recapitulate the hierarchical organization of the shoot. Collectively, these findings indicate that somatic epimutations act as a developmental precursor to heritable epigenetic variation and provide further support for a SAM-origin model of spontaneous epimutation formation.

## Methods and Materials

### Plant material

Transgenerational data: We used bulk whole-genome bisulfite sequencing (WGBS) data of single leaves collected from individual plants at different generations of the MA3 mutation accumulation pedigree, as presented in Shahryary et al. (**Fig. 1C**) (14). We used this material to quantify DNA heterogeneity within single leaves of individual plants and to study epimutation accumulation across generations.

Intra-plant data: Col-0 seeds were planted and grown for 5 weeks. Multiple leaves from a single plant were sampled and flash frozen in liquid nitrogen. DNA was isolated using a Qiagen Plant DNeasy kit (Qiagen, Valencia, CA, USA) according to the manufacturer’s instructions.

### Library preparation and sequencing

For the intra-plant data, WGBS libraries were prepared according to the protocol described in Zhou et al. (20). Libraries were sequenced on a NovaSeq X (Illumina) in a paired-end 150 bp format.

### Data processing

All WGBS data were processed with the MethylStar pipeline v1.4 (30). In particular, sequencing quality was assessed with FastQC v0.12.1 (31), and read trimming was performed with Trimmomatic v0.39.2 (32) using the parameter: ILLUMINACLIP:TruSeq3-PE.fa:1:30:9 LEADING:20 TRAILING:20, SLIDINGWINDOW: 4:20 MINLEN:36. The clean reads were aligned to the reference genome downloaded from Ensembl Plants (https://ftp.ensemblgenomes.ebi.ac.uk/pub/plants), removed duplicates, and extracted the methylation levels using Bismark v0.24.1 with default parameters (33). METHimpute v1.24 was used for cytosine methylation state calling (34). To ensure high-quality methylation state calls, cytosines with a maximum posterior probability larger than 0.99 (posteriorMax) were retained for downstream analyses.

### Annotation

We considered the following annotations: genes, transposable elements (TEs), promoters, gene-body methylated (gbM) genes, and sparsely methylated regions within “red” chromatin states (redCS-SPMRs). Gene annotations were downloaded from Ensembl Plants (https://ftp.ensemblgenomes.ebi.ac.uk/pub/plants), release 57. TE annotations were downloaded from The Arabidopsis Information Resource (TAIR) (35), release TAIR10. Promoters were defined as 1 kb upstream from the transcription start site of genes, downloaded from TAIR (TAIR10_upstream_1000_20101104). GbM genes and lowly methylated (LM) genes were retrieved from Zhang et al. (36). Note that the study classified LM genes as unmethylated genes to clearly distinguish them from gbM genes (36). However, Zhang et al. (37) showed that these genes feature low steady-state CG methylation, on average, rather than being strictly unmethylated. RedCS-SPMRs coordinates, which are defined by the intersection of a specific cluster of “red” Chromatin States (specifically CS 4, CS 5, and CS 6, highly enriched in gbM genes) and Sparsely Methylated Regions (SPMRs, characterized by stable, intermediate methylation levels ranging between 20% and 80%), were obtained from Hazarika et al. (8). The list of 24-nucleotide small RNAs (sRNAs), as known as the RNA-directed DNA methylation (RdDM) targets, was obtained from Slotkin et al. (unpublished data). Because of overlapping annotations, some cytosines corresponded to multiple annotations. To overlap methylomes with annotations, we used bedtools v2.31.1 (38).

### Quantifying CG methylation heterogeneity

To quantify heterogeneity in the CG context, we used the quantitative Fraction of Discordant Read Pairs (qFDRP) that is defined as the normalized Hamming distance of pairwise reads that cover individual CG sites (28) (**Fig. 1B**). The qFDRP score best captures the stochastic gains and losses of mCG characteristic of spontaneous epimutations occurring across the distinct cell layers (28) among the available read-discordance metrics (24–29) (**Fig. 1B**). We utilized the Metheor software v0.1.8 (29) to identify qFDRP site coordinates and scores with the command: metheor qfdrp --input Sample.bam --output Sample.tsv --min-qual 10 --min-depth 6 --max-depth 40 --min-overlap 35. The output coordinates are in a 0-based system. CG sites with qFDRP > 0 were defined as heterogeneous sites.

### Enrichment analysis

To assess the enrichment of heterogeneous sites (qFDRP score > 0) per annotation, we compared the observed (*O*) number of heterogeneous sites in each annotation to the expected (*E*) number based on how many CG sites the annotation contains. For a feature *f*, we defined *O*_*f*_ as the count of heterogeneous sites overlapping *f*, and *N*_*f*_ as the count of CG sites within *f*. We estimated the background heterogeneous rate *(p*) across the analysis universe (*U*) (such as the genome-wide or within a specific annotation) as *p*_*o*_ *= O*_*U*_ */ N*_*U*_, where *O*_*U*_ is the total number of heterogeneous sites in the background and *N*_*U*_ is the total number of CGs in the same background. And the expected count for a feature *f* was then computed as *E*_*f*_ *= N*_*f*_ *\* p*_*o*_. Enrichment was calculated as *Enrichment*_*f*_ *= O*_*f*_ */ E*_*f*_, where values > 1 indicate an excess of heterogeneous sites relative to expectation and values < 1 indicate depletion.

### Epimutation rate estimation

We estimated global and annotation-specific CG methylation gain (α) and loss (β) rates using the R/Bioconductor package AlphaBeta (v1.16.0) (14). For the transgenerational dataset (the MA3 pedigree), we used the heterogeneous site coordinates identified in generation 1, line 2 (G1L2) as a reference, and then retrieved the methylation levels and heterogeneity scores at those sites in other generations. Next, we fit the ABneutral model to mCG divergence data, with divergence time expressed in generations; this yielded CG epimutation rates per site, per haploid genome, per generation (5,8,14,39). For intra-plant analyses, we used ABneutralSOMA. As in our previous tree-based work (14,19,20), we represented plant branching architecture as a pedigree of somatic cell lineages, with sampled leaves as lineage endpoints. Leaf-to-leaf divergence was quantified as normalized physical distance (centimeter), so epimutation rates are reported per site, per haploid genome, per unit of normalized physical distance.

### Gene ontology analysis

Gene Ontology (GO) analysis (GO biological process) was performed using Protein ANalysis THrough Evolutionary Relationships (PANTHER) v19.0 (http://www.pantherdb.org/) (40).

## Results

### Somatic CG methylation heterogeneity in single leaves

To quantify somatic CG methylation (mCG) heterogeneity, we reanalyzed WGBS data from single leaves of two separate *A. thaliana* plants. The two plants were generation 1 (G1) individuals sampled from separate lineages of the MA3 pedigree (14) (**Fig. 1C**). The MA3 pedigree was originally generated by repeated selfing and single-seed descent using a progenitor design, in which the same plant that contributed the seed for the next generation was also sampled for WGBS (3,41). Because each generation passes through a strict single-cell bottleneck at seed formation, any cell-to-cell mCG variation observed within a sampled leaf must have arisen *de novo* during that plant’s own development. In a SAM-origin model, a subset of such changes will be stochastically incorporated into the reproductive lineage and accumulate across MA lines through epigenetic drift (4). Together, this experimental system provided a framework for investigating how somatic mCG heterogeneity within leaves relates to the transgenerational epimutational processes driving long-term methylation divergence.

We first quantified the extent and genomic distribution of somatic mCG heterogeneity using the quantitative Fraction of Discordant Read Pairs (qFDRP) metric. Metheor (29) was used to compute per-CG heterogeneity scores within each leaf. qFDRP scores ranged from 0 to 0.74, with higher values indicating greater read-level methylation discordance. Using the parameters (see **Methods**) and a liberal threshold of qFDRP > 0, we identified 2,056,810 heterogeneous sites across the genome (corresponding to ∼ 37% of all 5,567,714 CG sites, in both strands). Genome-wide qFDRP scores varied substantially (range: 0.0001778 - 0.74) yet showed relatively similar heterogeneity distributions between the two plants (**Fig. 2A**). Indeed, nearest-neighbor analysis revealed that the majority of heterogeneous sites detected in separate plants occurred at identical or very nearby positions (mean distance = 140 bp; median distance ∼ 0 bp) (**Fig. 2B**), suggesting that specific genomic regions are especially prone to stochastic CG methylation.

**Figure 2:**
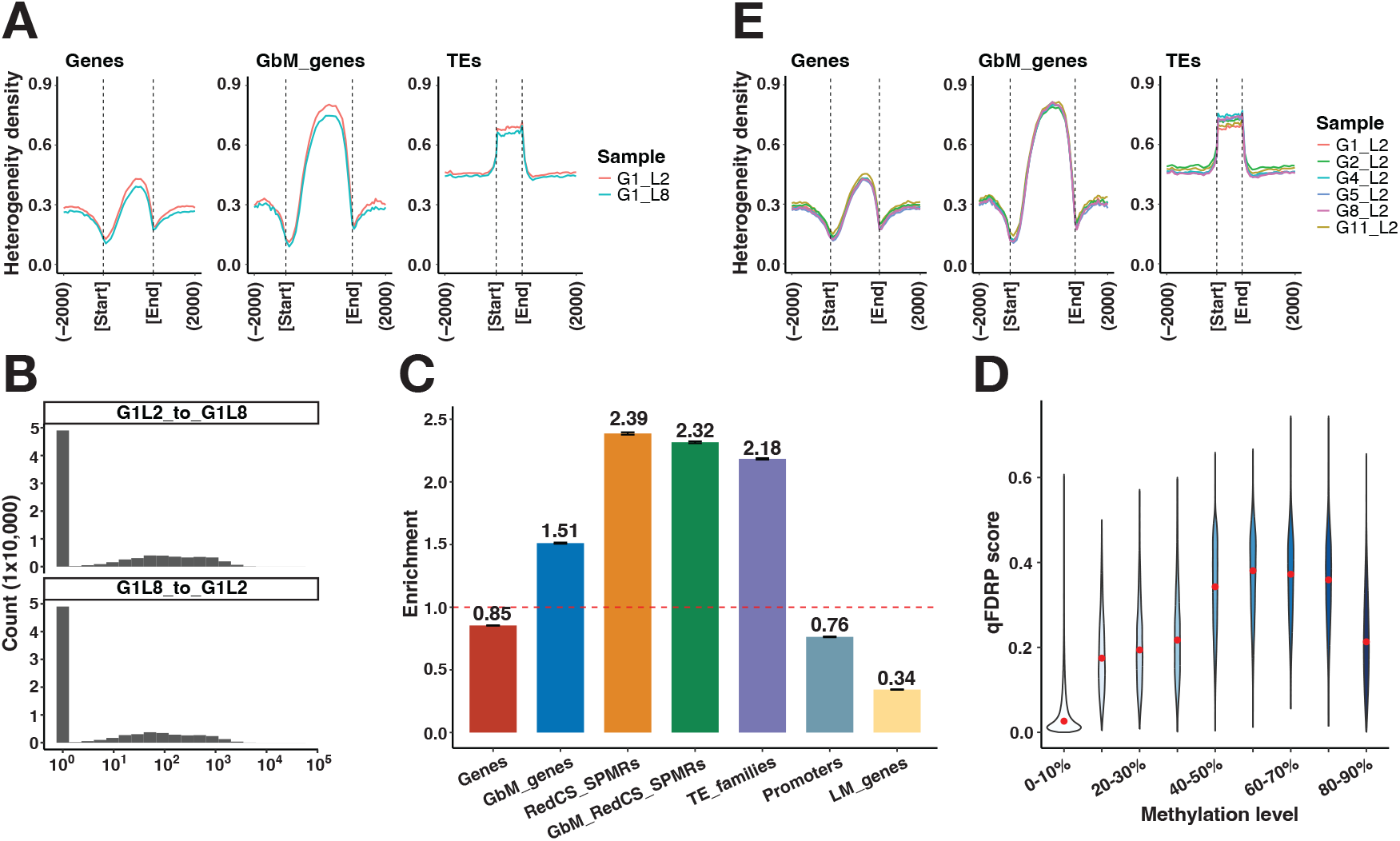
Genome-wide distribution, genomic enrichment, and temporal stability of heterogeneous sites in Arabidopsis. A. Spatial distribution of heterogeneous sites across genomic features including 2 kb up- and downstream, with dashed lines indicating feature start and end positions. B. Positional similarity of heterogeneous sites between independent plants was quantified as the distance from each heterogeneous site identified in one plant (e.g. G1L2) to the nearest heterogeneous site in the other plant (e.g. G1L8). C. Genomic enrichment of heterogeneous sites. The dashed line indicates the expected value (enrichment = 1). 1) Error bars represent 95% bootstrap confidence intervals, calculated with 1000 samples. D. Relationship between CG methylation level and qFDRP score. E. Stability of heterogeneity patterns across generations in the MA3 pedigree. GbM_RedCS_SPMRs: the intersection of gbM genes and redCS-SPMR regions.

Annotation analysis revealed that heterogeneous sites were significantly enriched in transposable elements (TEs) and gene-body methylated (gbM) genes, a conserved class of moderately expressed housekeeping genes characterized by high CG methylation and little or no non-CG methylation (42), with 2.18 and 1.51 fold enrichment, respectively (p = 0.0009, FDR ≤ 0.05, two-sided permutation test) (**Fig. 2C**). Because both gbM genes and TEs are strongly methylated in the CG context, their enrichment raises the possibility that heterogeneity might simply reflect absolute methylation levels. To test this, we examined median qFDRP scores across bins of mCG levels. qFDRP scores did not increase in a simple linear or monotonic way with methylation level (**Fig. 2D**). Instead, the relationship was strongly non-linear, and qFDRP scores also varied substantially within individual mCG bins. These results suggest that the heterogeneity scores capture specific instability in methylation maintenance rather than the methylation level itself.

In addition to the two G1 plants, we also analyzed single-leaf WGBS data from later-generation MA3 individuals (Line 2, G2 - G11). Their heterogeneity distributions shared the same trend with those observed in G1, and we detected no clear directional shift in genome-wide heterogeneity with generation (**Fig. 2E**). This temporal stability indicates that there is no cumulative transgenerational carryover of heterogeneity and supports the notion that it primarily captures *de novo*, within-generation somatic mCG variation established during individual plant development.

### Somatic heterogeneity converges on intragenic transgenerational epimutation hotspots

Heterogeneous sites in gbM genes were particularly interesting because this class of genes is known to harbor transgenerational CG epimutation hotspots in *A. thaliana* (8,39), and have recently been used as an epigenetic clock for evolutionary phylogenetic inference (5). We recently demonstrated that these hotspots are not uniformly distributed within gbM genes but instead correspond to subregions defined by specific chromatin states and sparsely methylated regions, which we have termed “redCS-SPMRs” (8) (see **Methods**). Although redCS-SPMRs comprise only ∼ 12% of all CG sites, they contribute ∼63% of CG epimutation events per generation in MA pedigrees (8). Interestingly, heterogeneous sites were strongly enriched in redCS-SPMRs relative to background gbM regions (2.32-fold vs. 1.51-fold enrichment, *p* = 0.0009, FDR ≤ 0.05, 1000 two-sided permutation test) (**Fig. 2C**).

The single-nucleotide resolution of qFDRP scores enabled us to examine transgenerational epimutational dynamics specifically at somatically heterogeneous CG sites. To do this, we used all samples (N = 13) in the MA3 pedigree and examined epimutation accumulation among lineages over 11 sexual generations (G1 - G11) (**Fig. 1C**). We found that mCG divergence among MA lineages increased substantially faster at heterogeneous sites in redCS-SPMRs within gbM regions than across all CGs genome-wide or across the broader sets of gbM or redCS-SPMR sites (**Fig. 3A**), consistent with their substantially elevated epimutation rates (**Fig. 3B**). Thus, heterogeneous sites detected in leaves not only predict epimutation hotspots identified in earlier multigenerational studies (8), but also pinpoint particularly labile sites nested within the broader hotspot domains defined previously.

**Figure 3:**
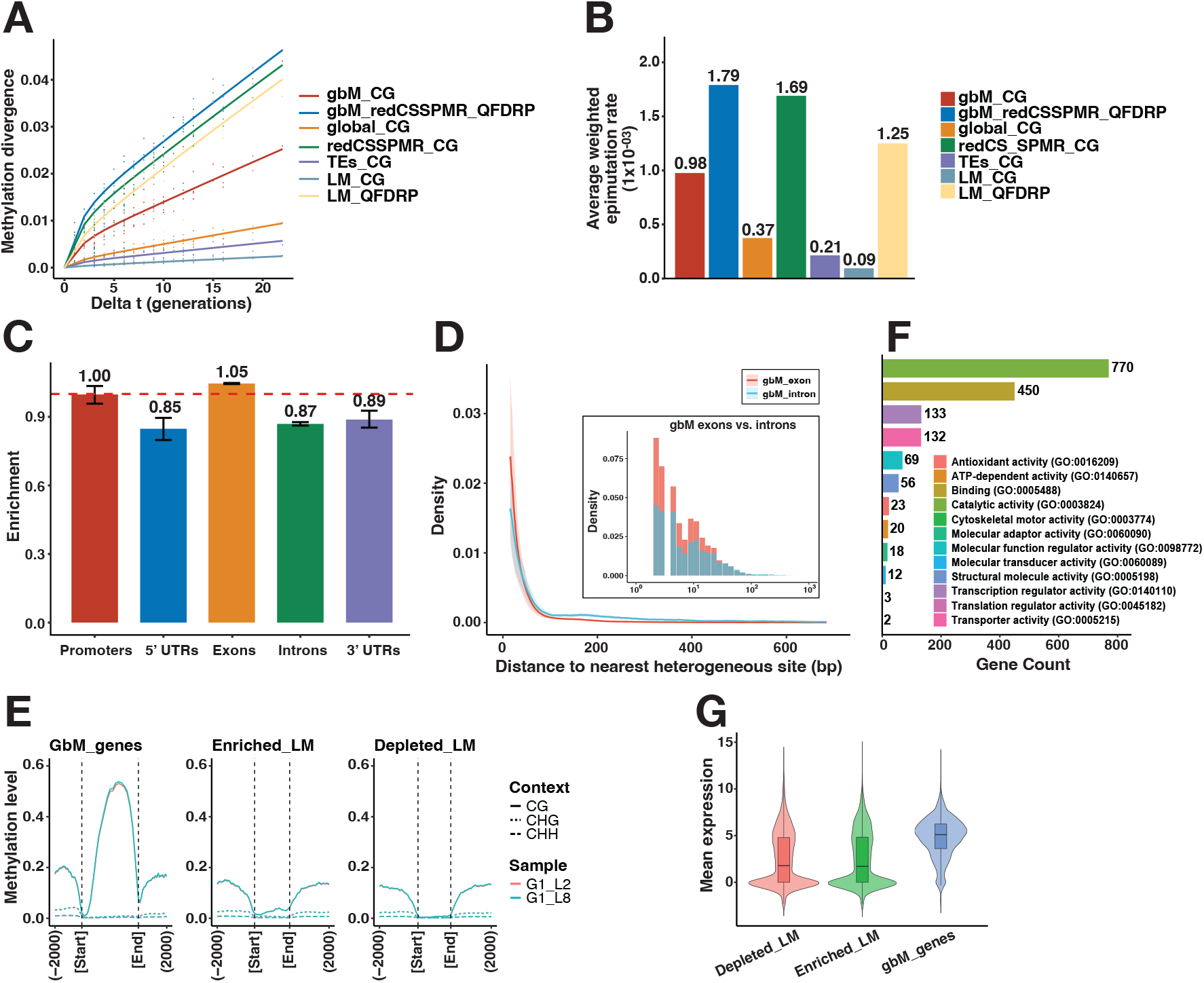
Heterogeneous sites identify transgenerational epimutation hotspots and reveal distinct classes of labile CG loci in Arabidopsis. A. Methylation divergence at heterogeneous (qFDRP score > 0) sites across generations. The fitted lines reflect methylation divergence (y-axis) as a function of divergence time (in generations; x-axis). B. Average weighted epimutation rates at heterogeneous sites across generations. C. Distribution of heterogeneous sites within gbM genes. The dashed line indicates the expected value (enrichment = 1). Error bars represent 95% bootstrap confidence intervals, calculated with 1000 samples. D. Spatial autocorrelation of heterogeneous sites within gbM genes, with the light-blue (introns) and pink (exons) bands corresponding to ± 1 standard deviation. Insets show the corresponding distributions of nearest heterogeneous site distances on a log10-scaled x-axis. E. Methylation profiles of heterogeneous-enriched (Enriched_LM) and -depleted (Depleted_LM) LM genes. F. Functional enrichment of Enriched_LM genes. G. The mean expression of LM gene subclasses. global_/gbM_/TEs_/LM_/redCSSPMR_CG: CG sites in the genome-wide, gbM genes, TEs, LM genes, and redCS-SPMRs; gbM_redCSSPMR_/LM_QFDPR: heterogeneous sites in the intersection of gbM genes and reCS-SPMRs, and in LM genes.

We examined the distribution of heterogeneous sites within gbM genes in greater detail. This allowed us to refine the localization of hotspot regions beyond the coarser chromatin-state annotations used in their original definition (8). We found that heterogeneous sites were modestly enriched in exons (**Fig. 3C**), which suggests that the most labile sites are preferentially located in pre-methylated coding regions (10). Within exons, pairwise distances between heterogeneous sites followed a steep exponential decay distribution (**Fig. 3D**), indicating that somatic methylation heterogeneity is strongly autocorrelated over short genomic distances. By comparison, in introns, which have lower steady-state CG methylation levels (**SI Fig. 1**), pairwise distances between heterogeneous sites were not only longer overall, but also decayed more slowly as a function of inter-CG spacing (**Fig. 3D**). This pattern is reminiscent of the recently proposed cooperative model of CG methylation maintenance (8), in which the propensity of a given CG to change methylation state depends on both the distance to the nearest neighboring CG and the methylation status of that neighbor. The observed autocorrelation among heterogeneous sites also explains why region-level epimutation rates can approach single-site rates (39), because gains and losses often occur in short clusters of adjacent CGs rather than as fully independent events at isolated sites.

In addition to gbM genes, qFDRP scores also identified transgenerational epimutation hotspots within lowly methylated (LM) genes (37) (see **Methods**). We previously showed that these LM genes, as a class, exhibit transgenerational epimutation rates that are ∼10.4-fold lower than in gbM genes. Here, however, we found that qFDRP-marked CG sites within LM genes diverged substantially faster among MA3 lineages than CG sites in LM genes overall (**Fig. 3A**). Indeed, epimutation rate estimates at heterogeneous sites within LM genes were ∼ 13.9 times higher than that in the background of these genes (1.25.10^-03^ vs. 9.00.10^-05^ per CG per haploid genome per generation, respectively) (**Fig. 3B**). To explore if heterogeneous sites index a specific subset of LM genes, we divided these genes into heterogeneous-enriched and heterogeneous-depleted sets (**Methods**). We found that heterogeneous-enriched LM genes resemble gbM-like genes in their CG and non-CG methylation profiles, but with substantially lower absolute mCG levels (**Fig. 3E**). This pattern suggests that many of these genes may fall just below the gbM classification thresholds used in earlier studies (43). Consistent with this interpretation, GO analysis indicated that heterogeneous-enriched LM genes are broadly associated with core cellular functions, in line with the housekeeping roles often linked to gene-body methylation (**Fig. 3F**). However, this similarity did not extend to transcriptional patterns. At the transcriptional level, heterogeneous-enriched LM genes more closely resembled core LM genes rather than gbM genes (**Fig. 3G**), which indicates that they represent an intermediate class combining the expression status of lowly methylated genes with subtle gbM-like methylation features. This observation further suggests that genic mCG maintenance errors are independent of transcriptional activity (37).

Taken together, these findings show that the same CG sites exhibiting pronounced somatic methylation discordance within leaves of an individual plant are also those that stochastically gain or lose methylation across generations in sexually propagated lineages, providing a base-pair-resolution view of the most active epimutational hotspots in the Arabidopsis genome.

### Somatically heterogeneous TE sites separate into distinct transgenerational classes by RdDM status

Our analyses also uncovered that leaf heterogeneous sites were strongly enriched in TEs (**Fig. 2B**). This is notable because TEs have been identified as epimutation coldspots in multigenerational experiments (1,3,39). One explanation for these transgenerational coldspots is RNA-directed DNA methylation (RdDM) reinforcement of TEs during gametogenesis and embryogenesis (3), which resets somatically acquired CG methylation changes before transmission to the next generation (11,13). This model predicts that TE sites can show substantial somatic heterogeneity within leaves while still exhibiting limited long-term heritable divergence.

To test this directly, we partitioned TE-associated heterogeneous sites using RdDM target coordinates (courtesy of the Slotkin lab) into two classes: sites in RdDM-targeted TEs and sites in non-RdDM-targeted TEs. The two classes had similar mean qFDRP scores (**Fig. 4A**), indicating that RdDM targeting does not measurably affect the accumulation of somatic methylation heterogeneity in leaves. Their transgenerational behavior, however, differed markedly. Methylation divergence accumulated significantly faster at heterogeneous sites in non-RdDM-targeted TEs than in RdDM-targeted TEs (**Fig. 4B**), and the average weighted epimutation rate of the non-targeted class was approximately 4.29-fold higher than that of the targeted class (3.61.10^-04^ vs. 8.40.10^-05^ per CG per haploid genome per generation; **Fig. 4C**). Thus, heterogeneous sites within TEs that fall outside RdDM target regions represent a relatively labile component of repetitive DNA. By contrast, heterogeneous sites within RdDM target regions show substantially greater transgenerational methylation fidelity. The same pattern was observed across major TE superfamilies (DNA/MuDR, LTR/Gypsy, and RC/Helitron), with non-targeted heterogeneous sites consistently showing faster divergence and higher weighted epimutation rates (**SI Fig. 2A, B**). We also examined the spatial distribution of non-RdDM-targeted heterogeneous sites within TE bodies to test whether epimutation-prone sites cluster in specific TE subregions, but found no clear positional bias (**SI Fig. 2C**). These results help explain why previous estimates of mCG epimutation rates in TEs, which pooled RdDM-targeted and non-targeted sites, yielded only composite TE-wide averages and could not distinguish these distinct transgenerational behaviors. By combining single-site heterogeneity with RdDM annotation, our analysis uncovers this previously hidden heterogeneity and reveals a finer-scale epimutation landscape within repetitive DNA.

**Figure 4:**
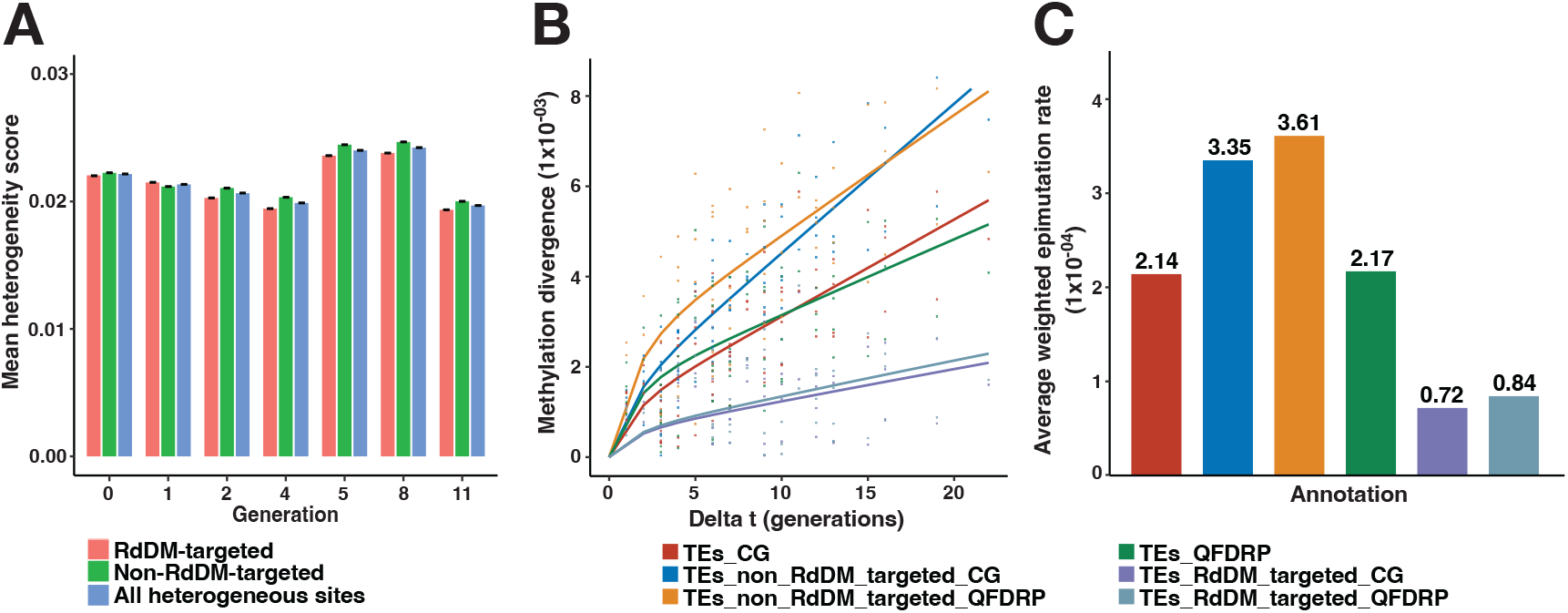
qFDRP identifies distinct transgenerational methylation stability related to the RdDM pathway at transposable element loci. A. Similar somatic methylation heterogeneity in RdDM-targeted and non-RdDM-targeted heterogeneous sites (qFDRP score > 0) in TEs. Error bars indicate standard errors from bootstrap. B. Methylation divergence across TE subclasses. The fitted lines reflect methylation divergence (y-axis) as a function of divergence time (in generations; x-axis). C. Average weighted epimutation rates across TE subclasses. TEs_CG/QFDRP: CG/heterogeneous sites in TEs; TEs_non_RdDM_targeted_CG/QFDRP: CG/heterogeneous sites in non-RdDM-targeted TEs; TEs_RdDM_targeted_CG/QFDRP: CG/heterogeneous sites in RdDM-targeted TEs.

### Intra-plant methylation divergence recapitulates shoot developmental architecture

The results above are strongly consistent with a SAM-origin model of spontaneous epimutations, in which stochastic errors in DNA methylation maintenance in somatic tissues propagate through development and manifest as transgenerational instability. A further prediction is that somatic epimutations accumulate along developmental lineages, generating structured methylation divergence among organs within an individual. If qFDRP scores can identify sites where methylation changes arise during somatic growth in the apical meristem, then leaves that are more widely separated in developmental space or time should show greater divergence at these loci than leaves derived from more closely related meristem progenitors (4,17,44).

To test this prediction, we performed WGBS on multiple leaves from a single *A. thaliana* individual (**Fig. 5A, Methods**). For each leaf, we performed per-CG methylation state calling, classifying sites as methylated or unmethylated (**Methods**). In bulk tissue, this type of calling is expected to capture epimutational states that have become fixed, or nearly fixed, within the predominant cellular lineage contributing to the sample (23). In leaves, this lineage is expected to derive primarily from the L2 layer of the shoot apical meristem, whose cells are proportionally most abundant (45). Focusing only on status calls of heterogeneous sites identified above, we quantified pairwise methylation divergence and related it to leaf positions along the shoot axis (14). Divergence at heterogeneous sites increased with plastochron distance (normalized centimeter along branches; **Fig. 5B**), consistent with lineage-dependent accumulation of somatic methylation instability during development. Notably, unsupervised clustering based only on heterogeneous-site divergence reconstructed the expected branching topology (**Fig. 5C and SI Fig. 3**), indicating that somatic variation at these loci retains information about shoot developmental history. Although divergence increased with developmental spacing, the relationship exhibited greater scatter than observed in long-lived perennial species (14,20). This variability likely reflects that polyclonal axillary meristems can contribute distinct lineages to a single lateral shoot (21,22), and that the shallow branching hierarchy of *A. thaliana* (Col-0) provides too few nested cellular bottlenecks for SAM-derived epimutations to become clonally fixed within leaves through somatic drift (4,17). In addition, developmental distances were inferred from physical spacing rather than true lineage distance (for example, number of intervening cell divisions or developmental time), which may further obscure the quantitative relationship.

**Figure 5:**
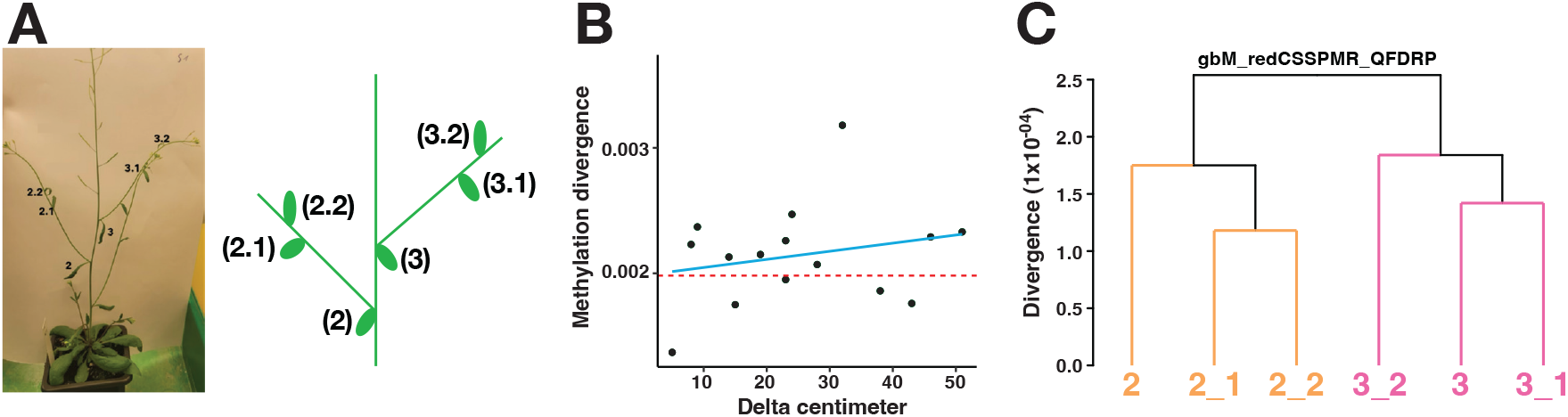
Somatic methylation divergence at heterogeneous sites reflects developmental lineage relationships within a single Arabidopsis plant. A. Sampling design for intra-plant methylation analysis. WGBS was performed on each leaf to obtain DNA methylation data. B. Pairwise methylation divergence at heterogeneous sites (y-axis) plotted against normalized branch distance (centimeters; x-axis). The light blue is the linear regression fit. The dashed line serves as a reference for no divergence. C. Hierarchical clustering of leaf samples based on pairwise methylation divergence at heterogeneous (qFDRP score > 0) sites within the intersection of gbM genes and redCS-SPMR regions recapitulates within-plant branching topology.

Despite these sources of biological noise, the observed spatial structuring of methylation divergence supports the hypothesis that heterogeneous sites retain meristem-derived lineage information during organ formation, further reinforcing the conclusion that somatic epimutation dynamics serve as a developmental precursor to heritable epigenetic variation.

## Discussion

By quantifying read-level CG methylation discordance within individual Arabidopsis leaves, we show that substantial somatic methylation heterogeneity exists within a single organ and must arise *de novo* during that plant’s development in the MA design. Importantly, qFDRP was not a trivial proxy for CG methylation level; although heterogeneous sites were enriched in highly methylated compartments, qFDRP values varied widely among loci with similar methylation levels, indicating localized differences in maintenance fidelity rather than methylation abundance per se.

A central result is the strong correspondence between somatically heterogeneous CGs and transgenerational epimutation hotspots mapped in the MA3 pedigree. This convergence is most pronounced within gbM genes, where heterogeneous sites are particularly enriched in redCS-SPMR subregions and show elevated divergence and rate estimates relative to broader genomic or domain-level baselines. Moreover, the single-base resolution of qFDRP highlights fine-scale structure within gbM genes, with labile sites preferentially located in exons, consistent with subgenic heterogeneity in methylation stability.

The enrichment of heterogeneous sites in TEs further refines the view that TEs are epimutational “cold spots.” Our stratified analyses suggest that instability is concentrated in a subset of TE CGs and is modulated by RdDM: divergence accumulates more strongly at non-RdDM-targeted heterogeneous sites, whereas their counterparts in RdDM-targeted TEs show signatures consistent with efficient remethylation that counteract somatic losses. Thus, qFDRP-based mapping resolves epimutational susceptibility within repetitive DNA at a scale inaccessible when averaging across entire elements.

Moreover, intra-plant comparisons provide an orthogonal test of the SAM-origin model. Methylation divergence at heterogeneous sites increased with plastochron distance, and divergence-based clustering reconstructed the expected shoot branching topology. The overall structuring supports lineage-dependent accumulation of somatic epimutations during development.

Together, these results unify somatic methylation heterogeneity, transgenerational epimutation hotspots, and developmental lineage structure in a single framework consistent with a meristematic origin of spontaneous epimutations. Our study demonstrates that read-level discordance from a single WGBS sample provides a practical route to identify loci intrinsically prone to methylation-maintenance errors, complementing multi-generational designs and enabling hotspot discovery without long-term pedigree experiments.

## Acknowledgements

This work was in part supported by the Bundesministerium für Bildung und Forschung (BTV, FJ) and the Deutsche Forschungsgemeinschaft (FJ).

---

**SI Figure 1:**
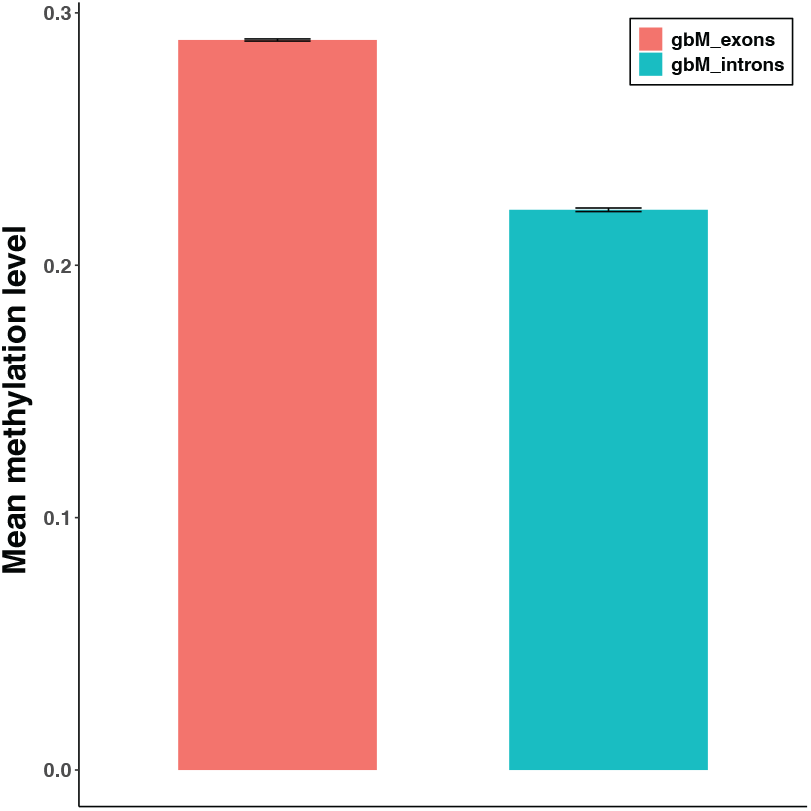
The mean expression levels in exons and introns within gbM genes. Error bars reflect standard errors from bootstrap.

**SI Figure 2:**
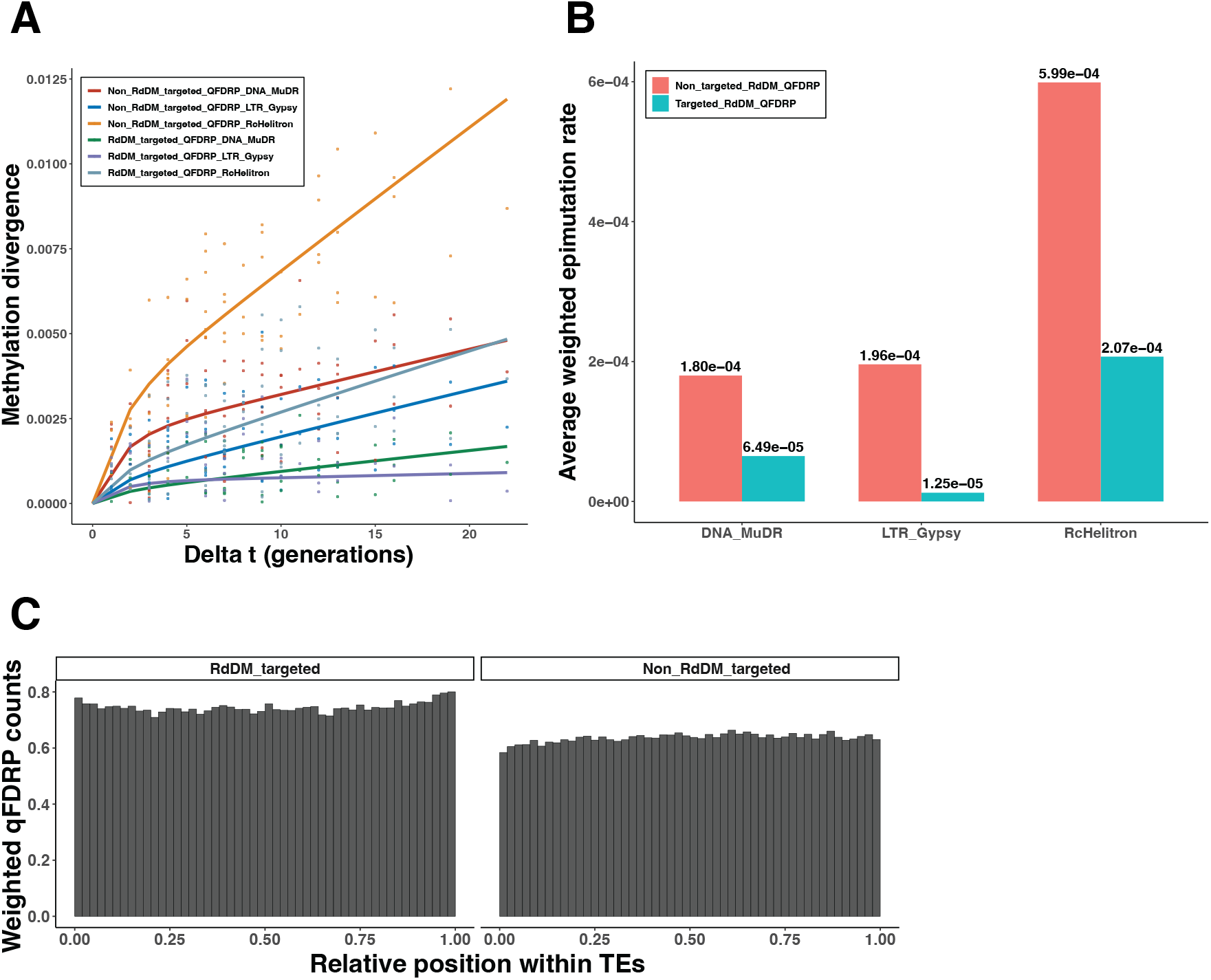
Methylation divergence and average weighted epimutation rates across TE super families. A. Methylation divergence at non-RdDM-targeted and RdDM-targeted heterogeneous (qFDRP score > 0) sites across TE superfamilies (DNA/MuDR, LTR/Gypsy, and RC/Helitron). The fitted lines reflect methylation divergence (y-axis) as a function of divergence time (in generations; x-axis). B. Average weighted epimutation rates at heterogeneous sites across TE super families. C. Spatial distribution of RdDM-targeted and non-RdDM-targeted heterogeneous sites within TE bodies. The heterogeneous site count is normalized by the CG count per bin (x-axis). The positions are calculated from the start position of each TE and normalized by TE length (y-axis). Non_RdDM_targeted_/RdDM_targeted_QFDRP: heterogeneous sites in non-RdDM-targeted and RdDM-targeted TEs.

**SI Figure 3:**
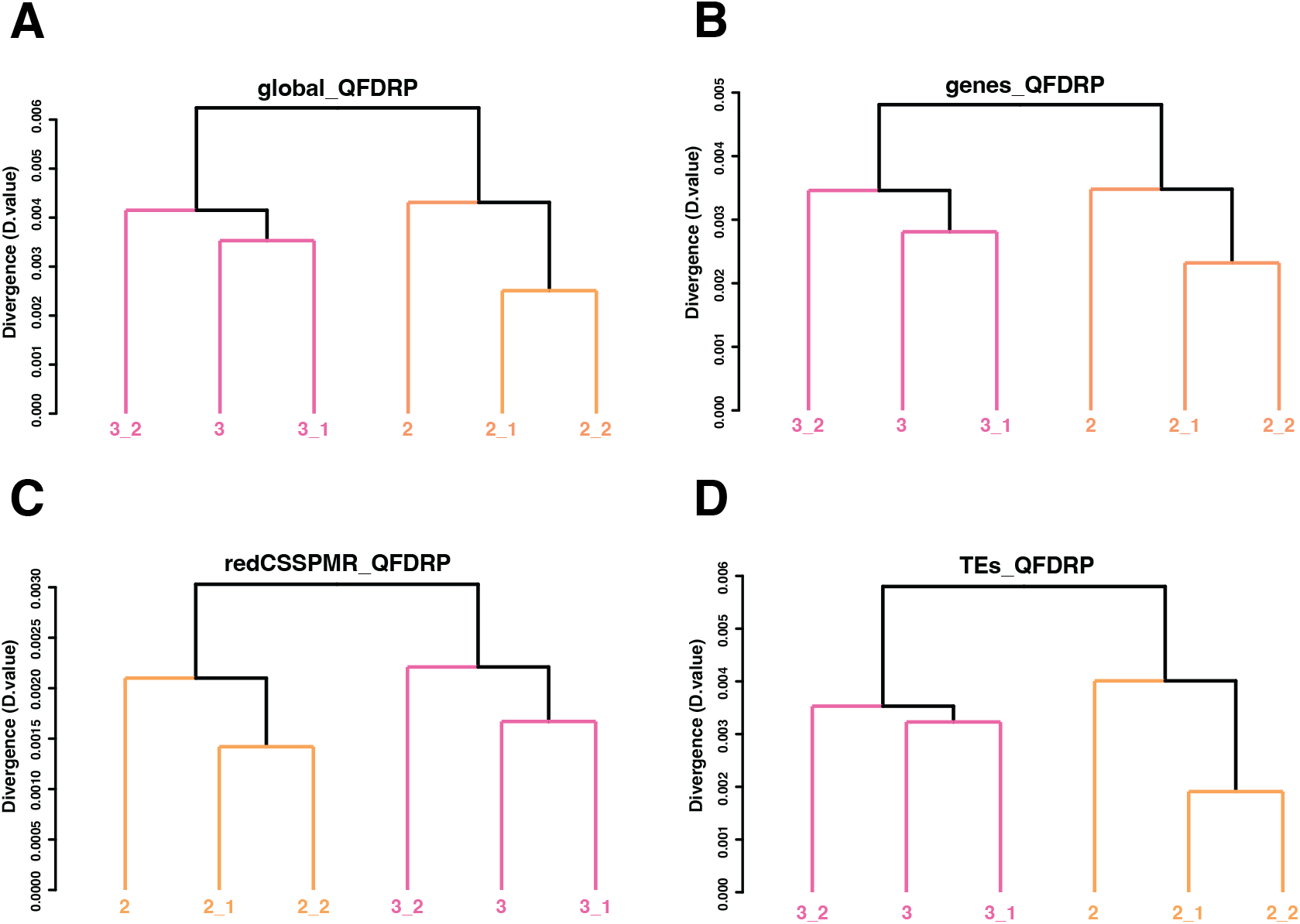
Hierarchical clustering of leaf samples based on pairwise methylation divergence at heterogeneous sites recapitulates within-plant branching topology. A. Using heterogeneous sites (qFDRP score > 0) in the genome-wide. B. Using heterogeneous sites in genes. C. Using heterogeneous sites in redCS-SPMRs. D. Using heterogeneous sites in TEs.

## Notes

### Competing Interest Statement

The authors have declared no competing interest.

